# Public perceptions, knowledge and factors associated with the acceptability of genetically modified foods in Kampala city, Uganda

**DOI:** 10.1101/2022.11.23.517645

**Authors:** Miria Nowamukama

## Abstract

Food is a basic human need important for the survival of all human beings. The rapidly growing world population puts pressure on food sources, inviting the need to devise mechanisms to sustain it. Biotechnology has developed important measures for improving plants and livestock as a remedy for food security with aim to achieve the Sustainable Development goal two (2). Globally, the adoption and use of genetically modified foods (GMFs) has been controversial as it is in Uganda today due to concerns related to the risk uncertainties. This study was therefore conducted to assess the public perceptions, knowledge and factors that influence the acceptability of genetically modified foods in Kampala City.

This was a cross sectional quantitative study that involved one hundred and ninety-eight participants. The data were collected using a survey tool and summarized using descriptive and linear regression analysis.

The findings of this study showed that almost two-thirds of participants (129/198, 65%) had some basic knowledge on genetically modified foods. About 45.3% (90/198) of the participants perceived genetically modified foods as being unsafe for human consumption. Eighty-eight participants (44.3%) perceived them as being associated with major human health and environmental safety concerns. The acceptability of these foods was significantly associated with gender, education level, nutritional value and health effects. Female participants were more likely to accept genetically modified foods (OR.4.84 95% CI: 1.37 - 7.68). Those who perceived genetically modified foods as being of high nutrition value were more likely to accept them (OR. 3.07, 95% CI: 1.27 - 7.37).

The public is predominantly aware of genetical modified foods since a big proportion had basic knowledge about them although with a lot of misinformation. People with a higher education level had positive perceptions on the use of these foods hence a need to educate the public to dispel misinformation that influences their acceptability

## Introduction

Hunger is one of the world’s greatest challenges that causes about 1.9 million deaths every year(1). Recently, there has been a global increase in the rates of malnutrition and undernutrition as resulting into exponential vulnerabilities of food insecurity due to climate change(2). The impact of climate change on the global hydrological cycle influences the patterns of supply of water for agriculture(3). This puts the poor at a much more risk of vulnerability since agriculture is the main source of their livelihood. In addition many communities have become disadvantaged due to lack of resilient systems and sufficient food reserves that can support the inhabiting populations(3).

The global fight against hunger as outlined in the 2030 sustainable development goals (SDG) agenda, remains a public health challenge in Low- and middle-income countries(4). The 2030 vision for zero hunger to promote healthy lives and thriving communities as stipulated in SDGs 2, 1 and 3 calls upon United Nations (UN) member states to sustainably work towards ending hunger and hunger-related diseases such as marasmus, kwashiorkor, anemia and goiter(5).

The emerging technologies have demonstrated the potential in reducing hunger and hence the contributed to improved strategies against the impact of climate change which in turn improve the individual and public health(6).

Biotechnology is one of these new technologies that has shown potential through its several applications in various sectors such as medicine, environment, and agriculture. The latter was of much interest since Genetically Modified Foods (GMFs) were the focus of this study given the importance of food to the survival and livelihood of human beings. Climate change and environmental pollution have been reported as major challenges of the last century globally(7). These have constrained food sources affecting the potential to sustain the rapidly growing population. Molecular genetics has increasingly become a potential and important technique in the enhancement of plant and animals for food production to ensure food security (8, 9).

However, the adoption of genetically modified foods has been controversial in many countries across the globe(10). The controversy mainly arises from the perceived risks and benefits of this technology(11). However, the ongoing public debates and controversy has been due to difference in risk benefit perception among different social groups in the public due to fear of the unknown and the perceived uncertainty of risk of this technology(12).

Genetically modified (GM) crops have gained acceptance in some countries although there has been increasing concerns on their safety for human health and environment which has raised several ethical concerns. This has resulted into significant differences among consumers’ perspectives regarding GM technology and its products across the globe(13).

Uganda currently has ongoing confined field trials since 2007 for GM crops such as bananas, maize, potatoes, rice and cassava though these have not yet been released on the Ugandan market(14). The previous attempt to pass the 2012 Biotechnology and Biosafety bill into law by the 10^th^Ugandan parliament on 4^th^October 2017 was unsuccessful. The Ugandan president declined to sign the bill mainly because it lacked clauses regarding protection of patent rights of indigenous farmers and sanctions for scientists who mix GMOs with indigenous crops and animals(15). He also pointed out that scientists and a few key experts were the only categories that had been consulted in the process and yet the public is a key stakeholder in this endeavor as consumers and end users. This study therefore aimed to explore public knowledge, perceptions and factors that influence the acceptability of genetically modified foods and by the end of the study, this was greatly achieved.

## Materials and methods

### Study area

This study was conducted within Kampala city to sample the Ugandan population because it was not possible to conduct a countrywide-study due to a limited funds. All the five divisions were stratified to obtain a representative sample from each division. These are Kampala Central, Nakawa, Makindye, Kawempe and Rubaga divisions. The study area was selected due its diversity as a capital city of Uganda. It’s composed of people from diverse cultures and religious affiliations to obtain an appropriate representation of the Ugandan population. The map below shows the location of Kampala city on the Ugandan map and displays the partitions and administrative boundaries of different divisions and the neighboring areas.

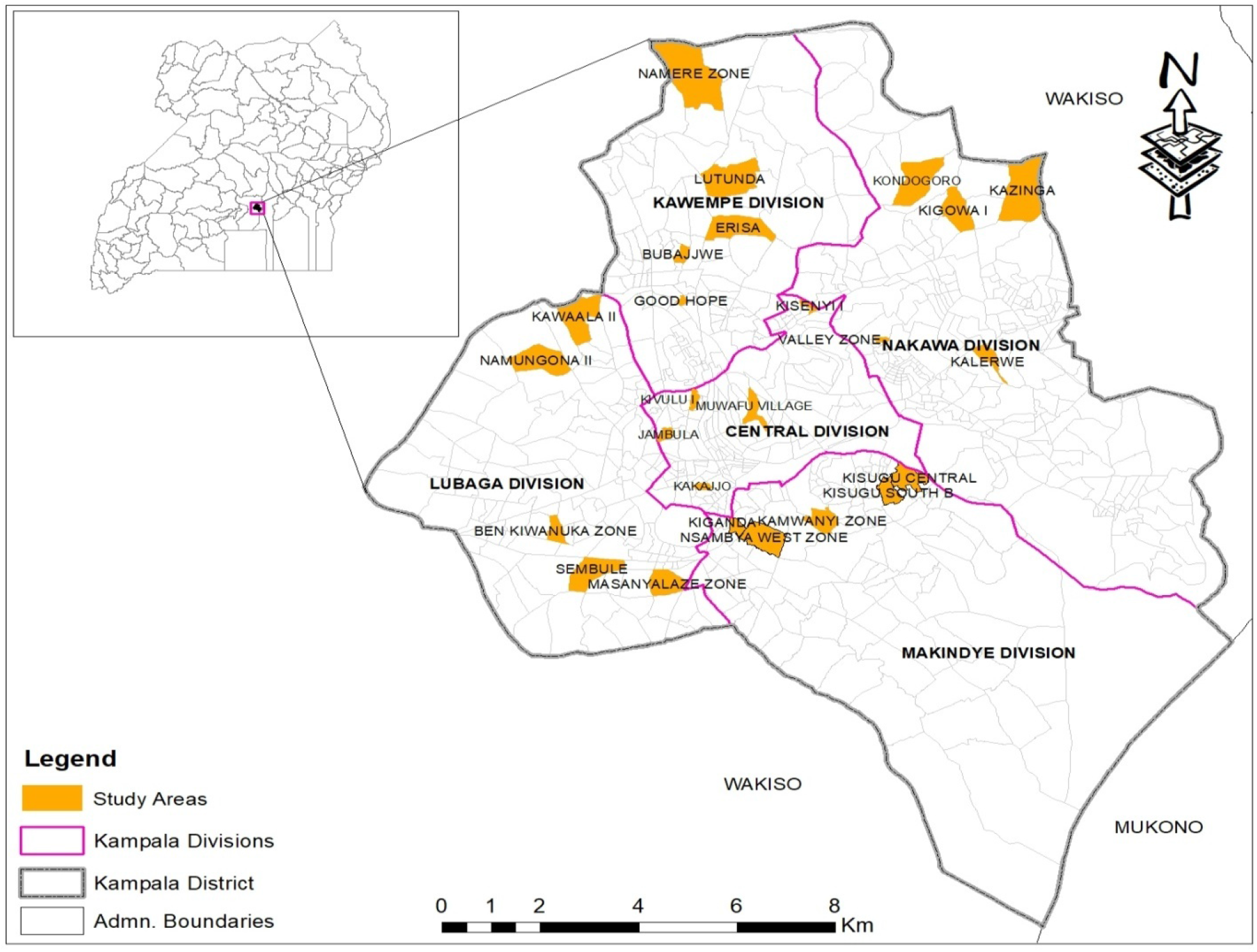

### Study design

This was a cross sectional quantitative design that involved the members of the public from varying backgrounds. One hundred ninety-eight individuals were selected across the five divisions of Kampala Capital City Authority (KCCA) to represent the public’s opinion. This survey recruited and consented adult individuals to participate in this study from May to July 2018.

### Data collection method

Data were collected by use of self-administered questionnaires which comprised of closed questions. The questions were categorized in four domains that included: demographic characteristics, basic knowledge on GMFs, perceptions, and factors that influence acceptability of GMFs in Kampala City. The Likert scale was used mainly for questions regarding acceptability to measure the level of agreement that ranged from strongly agree (1) to strongly disagree (5). Two graduate research assistants obtained 3 days training and clearly understood the questions and the objectives of this study and data was collected under close supervision.

### Sampling procedure

Stratified sampling was used to select the sampling-units that included divisions (strata) and within each stratum (division) a separate sample was obtained. Villages were randomly selected from the enumeration area units as defined by Uganda Bureau of Statistics (UBOS). Households were selected by systematic random sampling at an interval of five. The sampling frame that lists all villages per division was obtained from the 2014 National Housing and Population Census(16). Each stratum sample size was determined based on proportionate to population size.

The total population of Kampala district was stratified to obtain a sample size for each division. The sample was allocated to each division proportional to its population size across the 5 strata. Respondents were selected from within the random Enumeration area (PSUs) and only one individual was selected per household.

### Quality control

The data collected was double checked to ensure that all questionnaires had complete information. In a situation where the respondent failed to provide complete information required, that such a respondent was replaced, and the corresponding questionnaire would be discarded. Double entry of data was also done to ensure completeness.

### Data analysis

Data analysis was done using IBM SPSS statistics for windows. Descriptive statistics were used to summarize the demographic characteristics, level of knowledge, level of acceptability and public perceptions of GMFs. Categorical variables were reported as percentages and continuous variables were expressed as means ± Standard Deviation (SD). Linear multiple regression analysis was used to investigate the other factors associated with public acceptability of GMFs. A p-value of less than 0.05 was statistically significant. The outcome variable, acceptability of GMFs, was analyzed as a continuous variable and defined on a five-point Likert scale. The level of agreement ranged from strongly Agree (1) to Strongly Disagree (5).

## Results

The data was obtained with a high level of participation (100%) because GMFs debate was trending discussion in the different media platforms at the time the study was conducted since the Biotechnology and Biosafety Bill 2012 had just been tabled for discussion in parliament. As a result, the public perceptions in this study have been clearly presented with the pro and anti-biotechnology views.

### Social demographics of participants from the general public

Most participants (172/198, 86.9%) had at least attained secondary level education and males had a higher education level than females. Slightly more than half of participants (103/198, 52%) were married and most were gainfully employed as shown in Table 1 below.

**Table 1:**
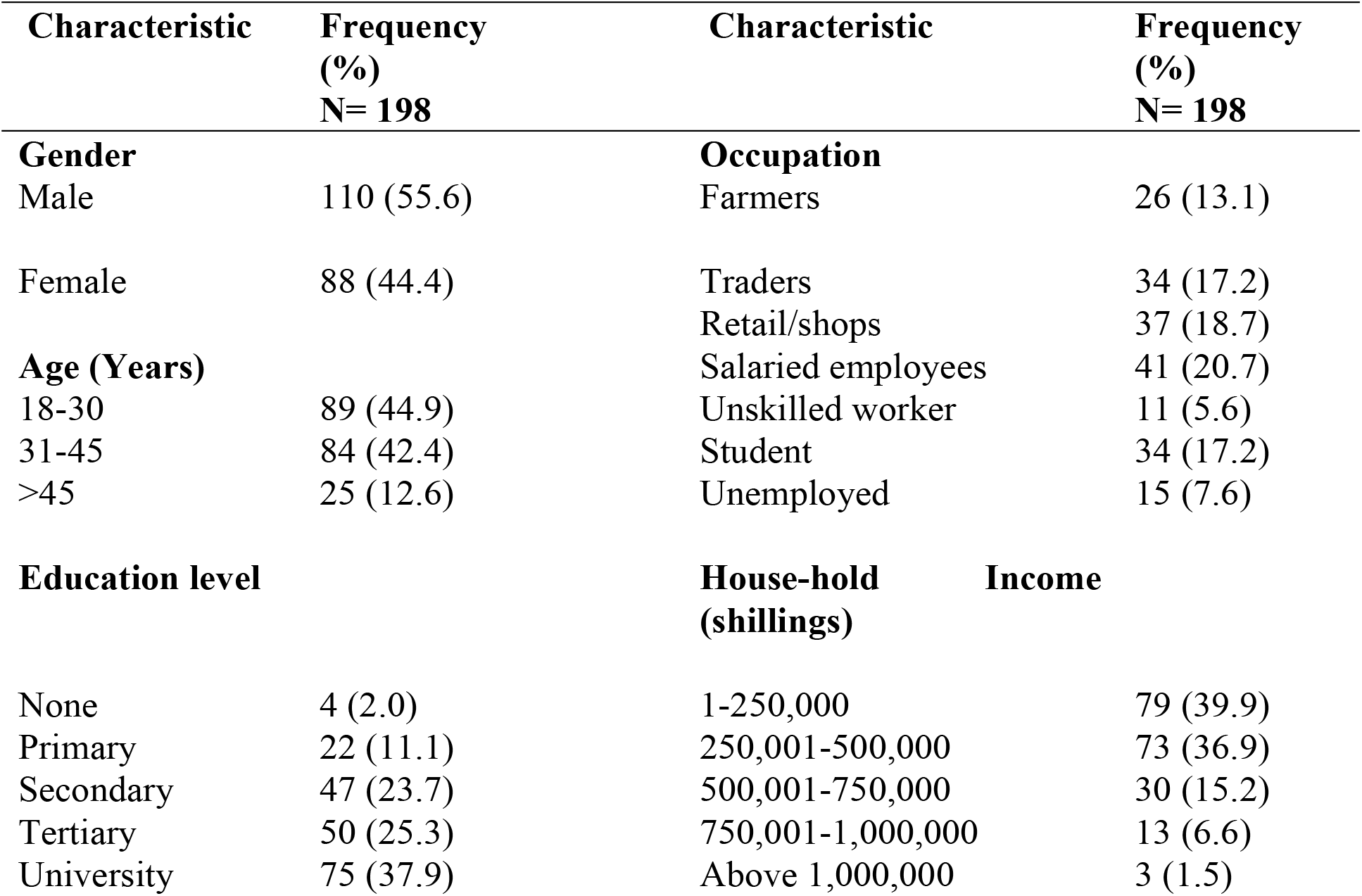
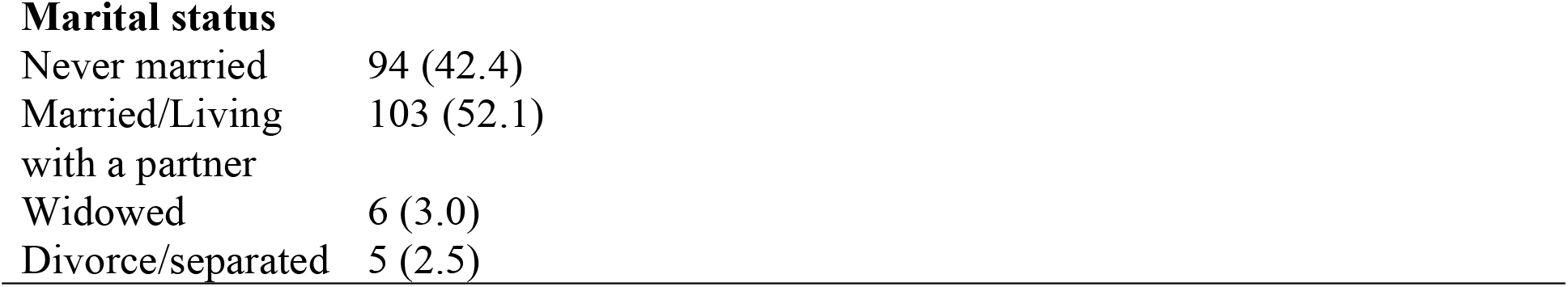
Social demographics of respondents

### Knowledge about Genetically Modified Foods

Almost two-thirds of participants (129/198, 65%) had ever heard about and had basic knowledge of GMFs. The main source of information on GMFs was from friends (121/198, 61%) and social media (56/198, 28%) followed by radio (46/198, 23%) and agricultural extension agents (46/198, 23%) as shown in Figure 1.

**Figure 1:**
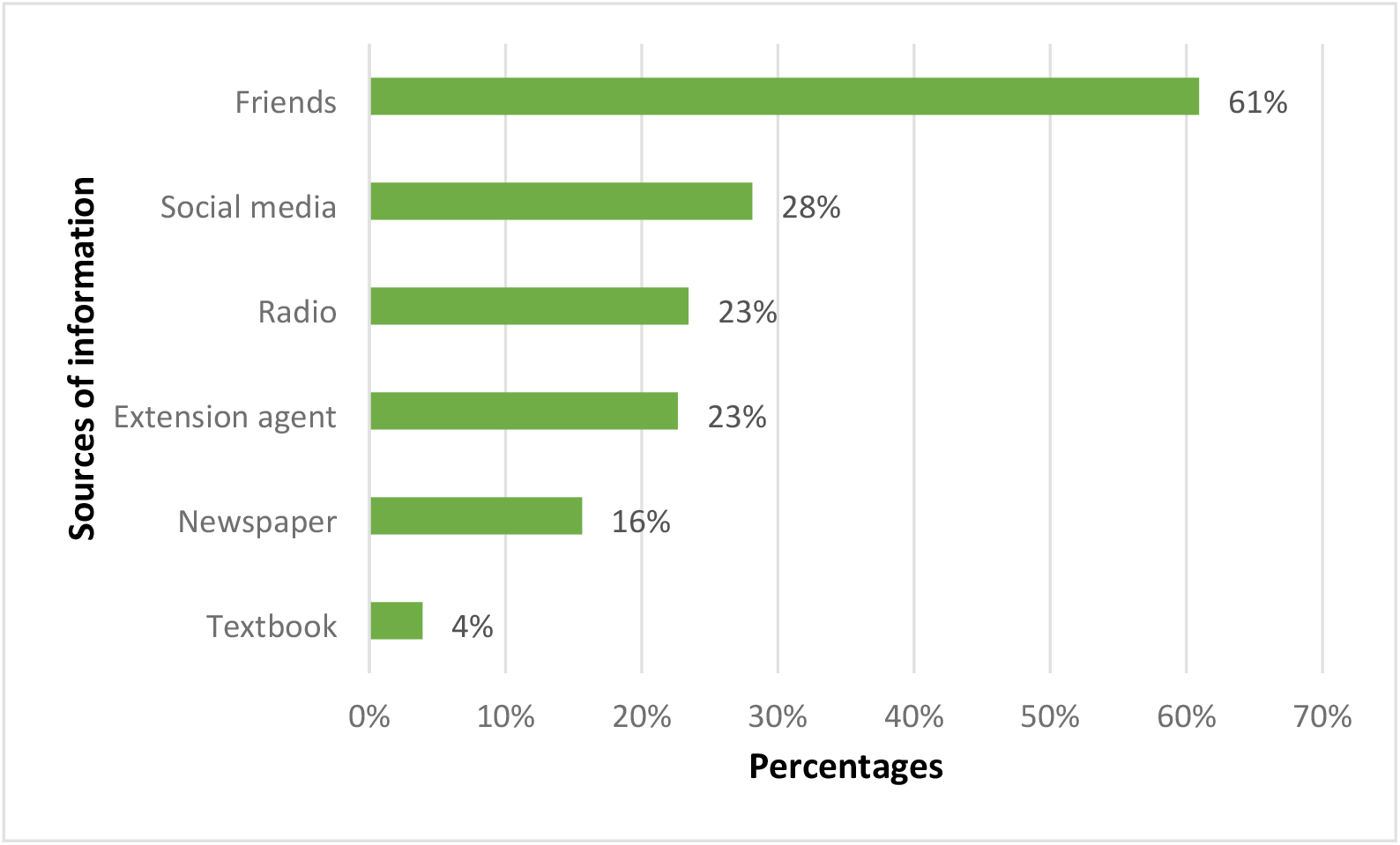
Sources of information on genetically modified foods

### Perceptions of risks about GMFs

Approximately forty six percent of participants perceived GMFs as being unsafe for human consumption (90/198, 45.3%) while (67/198,34.4%) were undecided about the safety of GMFs (Figure 2). Almost half of the participants (93/198, 47%) had other concerns about GMFs such as environmental and health effects.

**Figure 2:**
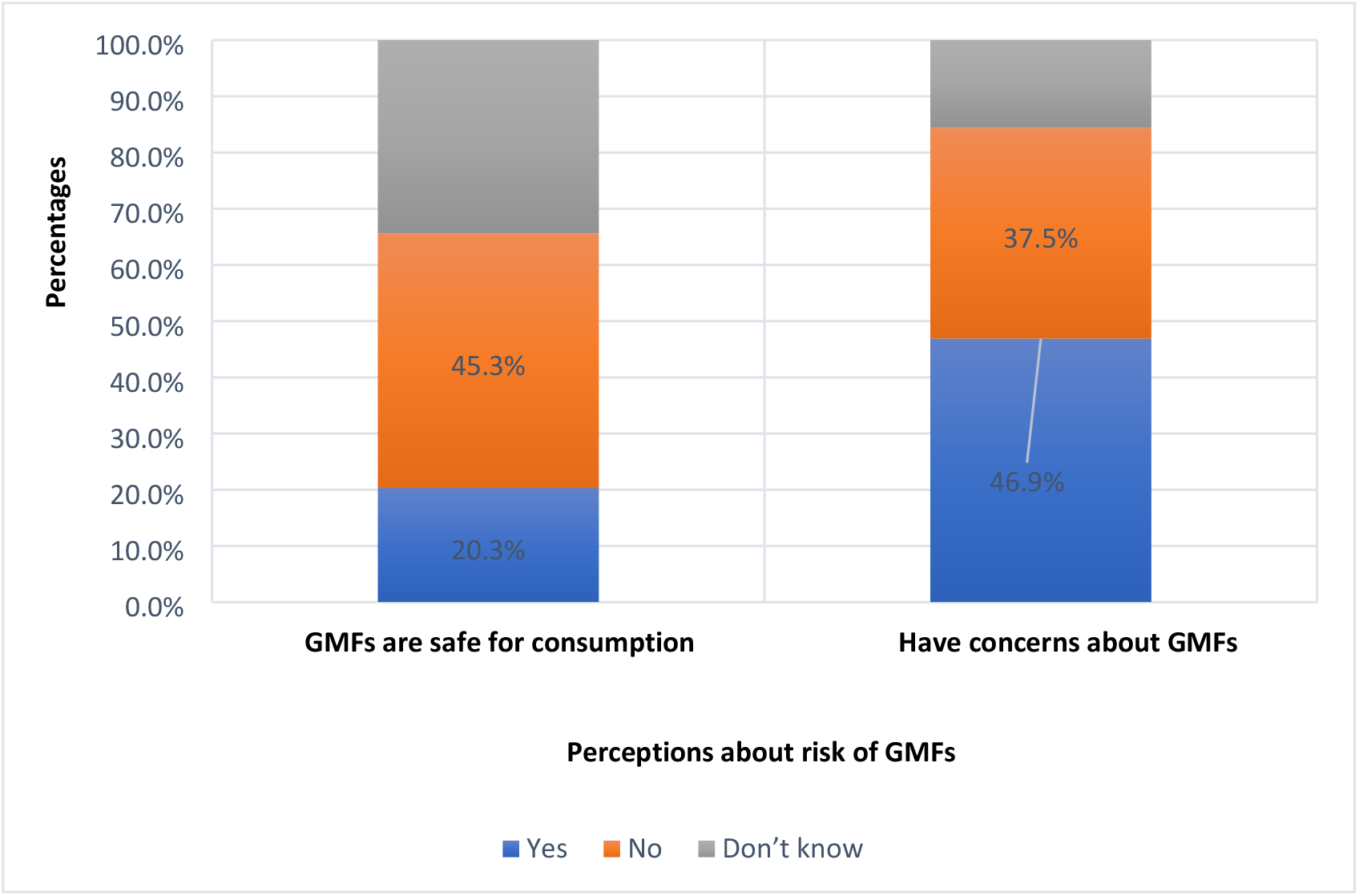
Perceptions about GMFs

**Figure 3:**
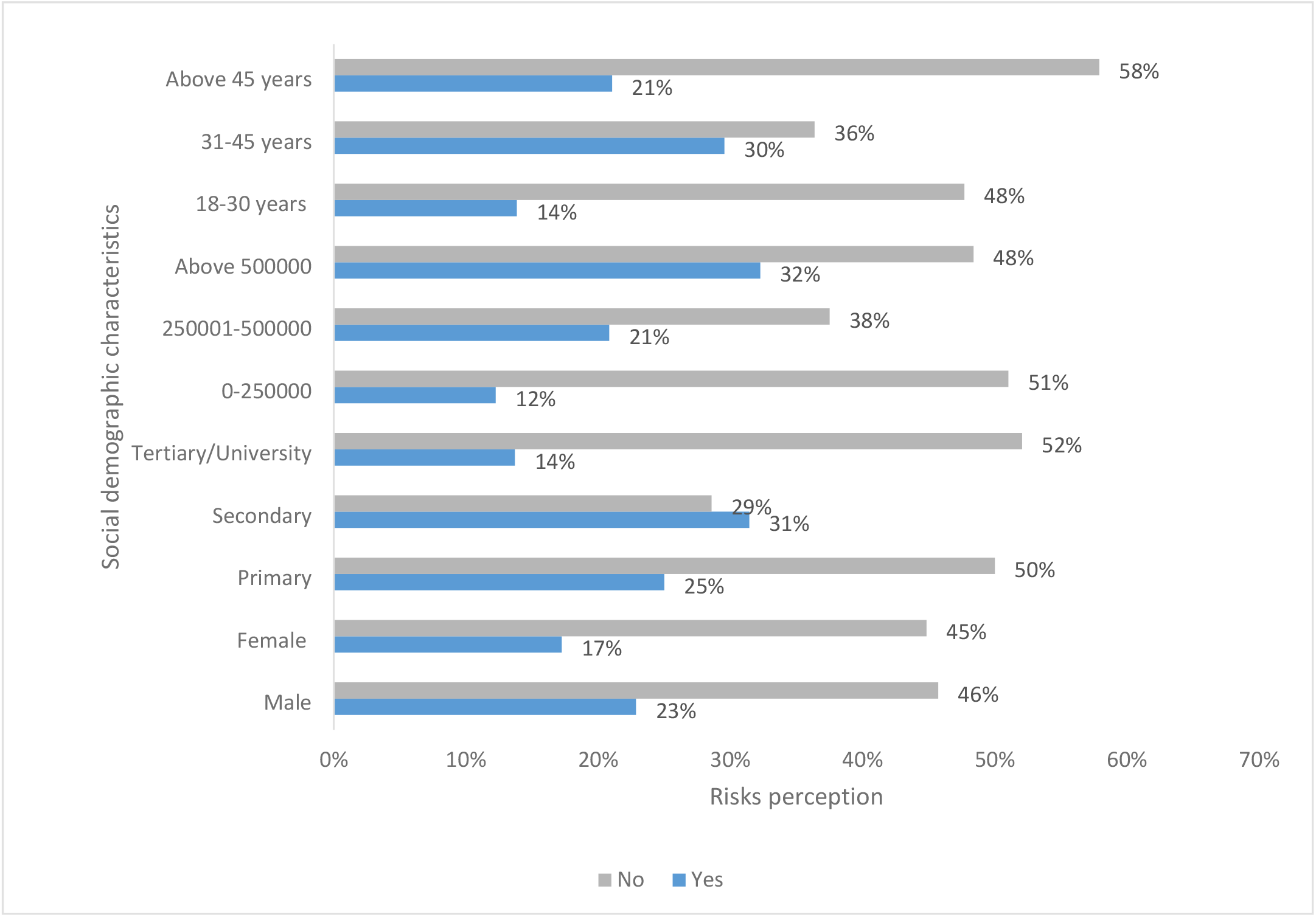
Participants’ perception of risk of genetically modified foods based on socio demographics Key; No; means that a participant perceived no risk on GMFs Yes; means that a participant perceived that GMFs are associated with risks.

Overall, individuals with ages ranging from 31 to 45 years (77%), females (149/198, 75%) and those with higher education levels agreed that GMFs are beneficial to humanity as summarized in Figure 4. The highly educated participants perceived GMFs as safe for human consumption and yet Individuals with just secondary level of education expressed more negative perception of GMFs due to the high risk perceived and other misconceptions.

Most concerns about GMFs were associated with risks to human health (88/198, 44.3%) and the environment (56/198, 28.4%). More than half (108/198, 54.5%) indicated that GMFs could tamper with nature

Female participants were more likely to accept GMFs compared to their male counterparts (OR. 1.99, 95% CI: 1.03-3.84, p= 0.04). Those who had attained tertiary level of education were three times more likely to accept GMFs compared to those who had never attained education or those that had less levels (OR. 3.01 95% CI: 1.09 - 9.51, p= 0.05). Similarly, those who perceived GMFs to be of high nutritive value were more likely to accept them as shown in Table 2.

**Table 2:** Factors associated with acceptability of GMFs.

| **Variable / Factor** | **Crude**<br>**OR** | **CI** | **P-value** | **Adjusted**<br>**OR** | **CI** | **P-value** |
| --- | --- | --- | --- | --- | --- | --- |
| **Age categories** |  |  |  |  |  |  |
| 18-34 | 1.00 | 1.00 | 1.00 | 1.00 | 1.00 | 1.00 |
| 35-45 | 0.81 | 0.41 - 1.62 | 0.81 | 0.16 | 0.02 - 1.69 | 0.13 |
| Above 45 | 0.43 | 0.15 - 1.22 | 0.43 | 0.11 | 0.01 - 6.81 | 0.3 |
| **Gender** |  |  |  |  |  |  |
| Male | 1.00 | 1.00 | 1.00 | 1.00 | 1.00 | 1.00 |
| Female | 1.99 | 1.03 - 3.84 | 0.04* | 4.84 | 1.37- 7.68 | 0.23 |
| **Marital status** |  |  |  |  |  |  |
| Never married | 1.00 | 1.00 | 1.00 | 1.00 | 1.00 | 1.00 |
| Married | 0.64 | 0.33 - 1.25 | 0.19 | 3.67 | 0.32 - 4.12 | 0.29 |
| Widowed/divorced | 0.29 | 1.57 - 1.53 | 0.15 | 0.65 | 0.009- 4.31 | 0.84 |
| **Religion** |  |  |  |  |  |  |
| Catholic | 1.00 | 1.00 | 1.00 | 1.00 | 1.00 | 1.00 |
| Protestant | 0.51 | 0.22 - 1.24 | 0.14 | 0.45 | 0.06 - 3.67 | 0.46 |
| Pentecostal | 0.64 | 0.26 - 1.52 | 0.31 | 0.65 | 0.04 - 1.3 | 0.78 |
| Moslem | 0.67 | 0.21 - 2.08 | 0.49 | 0.14 | 0.02 - 1.36 | 0.91 |
| Other | 0.59 | 0.17 - 2.12 | 0.42 | 0.03 | 0.06 - 1.49 | 0.08 |
| **Income** |  |  |  |  |  |  |
| Less than 250001 | 1.00 | 1.00 | 1.00 | 1.00 | 1.00 | 1.00 |
| 250001-500000 | 0.52 | 0.25 - 1.12 | 0.09 | 0.28 | 0.04 - 2.01 | 0.21 |
| Above 500000 | 0.99 | 0.44 - 2.25 | 0.99 | 2.07 | 0.24 - 17.97 | 0.51 |
| **Education level** |  |  |  |  |  |  |
| Primary/<br>Secondary | 1.00 | 1.00 | 1.00 | 1.00 | 1.00 | 1.00 |
| Tertiary | 3.01 | 1.09 - 9.51 | 0.05* | 1.33 | 0.36 - 4.27 | 0.08 |
| University | 1.44 | 0.51 - 4.11 | 0.49 | 0.71 | 0.29 - 17.42 | 0.83 |
| **Health effects** |  |  |  |  |  |  |
| No | 1.00 | 1.00 | 1.00 | 1.00 | 1.00 | 1.00 |
| Yes | 0.34 | 0.12 - 0.001 | 0.05* | 0.19 | 0.04 - 0.94 | 0.04* |
| **Not beneficial** |  |  |  |  |  |  |
| No | 1.00 | 1.00 | 1.00 | 1.00 | 1.00 | 1.00 |
| Yes | 0.25 | 0.53 - 1.21 | 0.08 | 0.07 | 0.001 - 4.15 | 0.2 |
| **Nutrition value** |  |  |  |  |  |  |
| Low | 1.00 | 1.00 | 1.00 | 1.00 | 1.00 | 1.00 |
| High | 3.46 | 1.53 - 7.86 | 0.003* | 3.07 | 1.27 - 7.37 | 0.01* |
| **Lower costs and taste** |  |  |  |  |  |  |
| No | 1.00 | 1.00 | 1.00 |  |  |  |
| Yes | 1.59 | 0.79 - 3.23 | 0.19 |  |  |  |
| **Social & political** |  |  |  |  |  |  |
| No | 1.00 | 1.00 | 1.00 |  |  |  |
| Yes | 1.05 | 0.23 - 4.84 | 0.96 |  |  |  |

\* Denote significant factor

When entered in a multivariate model only perceptions of high education levels, negative health effects and high nutritive value remained significant factors associated with GMFs acceptability as summarized in Table 2 above.

## Discussion

### Perception, knowledge, and acceptability towards GMFs

The application of biotechnology in the agricultural sector may be an important tool in Uganda since agriculture is a backbone of its economy. However, public concerns exist about its potential to negatively impact on the environment, human or animal health arising from different biotechnological processes and products(17). The findings in this study revealed that although majority of the residents in Kampala City are aware about GMFs, many seem to be unsure of their safety. Consequently, the high-risk perception among the study participants greatly influenced the relative lack of acceptability of these foods. Earlier studies have also reported similar findings alluding to high perceived risk of the GMFs and its low acceptability (18-20). Various policy interventions have been suggested and widely discussed in order to increase acceptability of GMFs especially among European consumers (20) although up to date, no consensus on evaluation of the safety of GMFs for consumption has been reached (18). Older studies also raised ethical concerns of the possibility that the technology used in GMFs could tamper with nature (21), something that has not yet been scientifically established up to date.

### Awareness about GMFs and associated risk

Two-thirds of the public reported having heard about GMFs. Like other earlier studies, public awareness on GMFs differs from the level of knowledge(22, 23). The findings in this study showed that majority of the public mainly obtained information from friends. However, friends may not be a reliable source of information because they may not necessarily be equipped with the technical knowledge regarding GMFs and this may perhaps explain the inadequacy in knowledge levels regarding GMFs by the public. Studies elsewhere have called for a need to have dependable sources of information regarding this controversial topic(13, 24-26).The public therefore requires adequate knowledge to make informed and independent decisions on GMFs. This will promote the principle of autonomy so that each individual can ably make their own dietary choice(27). The findings in this study are in agreement with other studies that consumer awareness influences the decisions and perceptions regarding the GMFs (13, 28).Therefore, the differences in public perceptions about GMFs ought to be respected since each individual is autonomous(29). However, for some cases, the social utility may override the personal autonomy in making dietary choices since the social benefit may be of a greater good to the majority than the individual choice(30). Most participants in this study believed that GM technology was associated with more risks than benefits and they also pointed out that this technology required adequate regulation that seemed to be a major limitation since there was no law on biosafety to govern its application at the time of this study.

### Factors affecting acceptability of genetically modified foods

Evidence has shown that although there is increasing knowledge on GMFs among the public, their acceptability is still wanting in many countries, and this was not any different in this study. The findings in this research revealed the significant factors that influenced the acceptability of genetically modified foods to be gender, education level, perceived risk or benefit, government trust, social and political issues.

The findings in this research revealed that, female participants were more likely to accept GMFs compared to their male counterparts. Women in the African settings are the core players in putting food on the table and engaging in agricultural production for a livelihood. Some earlier studies have also shown that women in developing countries play a big role in ensuring food security by holding its pillars such as its production, economy and nutritional security(31). Although on the contrary a study by Öz and colleagues revealed that males were more willing to accept consuming GMFs(29).

Furthermore, participants who had attained tertiary level of education were three times more likely to accept GMFs compared to those who had never attained education or those that had less levels. A study by Abedi has indicated that education has a great influence on public attitude towards genetically modified organisms(32). Therefore, the higher the level of education, the more the likelihood to accept to GMFs uptake or adoption.

Moreso, participants’ perception of risk or benefit greatly influenced their acceptability. The general perception may depend on the credibility of the source of the information and trust in the regulatory process(33). Relating to studies elsewhere, the perceived benefit had a positive impact on acceptability unlike the perceived risk (29, 34, 35). Similarly, findings in this research have shown that those who perceived GMFs to be of high nutritive value were more likely to accept them whereas those who highly perceived them as having adverse health effects were more likely to reject them(36).

The findings in this study also indicated that the public’ great fear about GMFs was the absence of a regulation. Another study conducted among Southern African Development Community (SADC) countries indicated that only a few of them in the region have biosafety frameworks for regulating the GM technology(37). Adequate regulation is paramount as a strategy to mitigate against any unanticipated risks and harms. Robust mitigation measures for protecting the environment and humans such as use of green houses, introduction of a trust fund, insurance covers and compensation ought to be introduced. Such fees should be paid by GM companies to the affected individuals and can be effectively implemented only if a law is in place. In addition, a law that incorporates the issues of environmental protection such as strict liability, appropriate labelling, collaborative partnership, and responsibilities of each body should be enforced to guard against any subsequent risks. Other studies emphasized the issue of labelling which could perhaps act as an appropriate market strategy so that a consumer may clearly make an informed decision and also as a measure to reduce the mixture of GM and non-GM food during harvesting or marketing(34, 38). The findings in this study further revealed that the public has less trust in government’s capacity to control, regulate and handle GMFs. The government may need to build trust through setting up policies and laws that will adequately protect its people and the environment in which they live. Scientists may need to contextualise the technology to the Ugandan setting, so that the science and innovation behind it is clearly understood by a local farmer and the consumer.

### Strengths and Limitations of the study

The study was conducted at a time when there was on-going discussion on the Biotechnology bill 2012 in the Ugandan Parliament which enhanced the relevancy of this study. However, some participants could hardly differentiate between the high breed food varieties from the GMFs and thereby mistook high breed varieties for GMFs. In addition, some household heads had a misconception that this study was a government initiative and hence misconceived that we were government spies. They suspected that their views would be disclosed as opponents to the GMO bill that was being discussed in parliament during that period. The study findings represent the public perceptions for the people in Kampala city and these may not be generalizable to the Ugandan population.

## Conclusion

Most members of the public had basic knowledge on GMFs. However, they had major concerns about the safety of GMFs to human health and the environment. Acceptability of GMFs was significantly influenced by female gender, high education level and the perceived nutritional value and health effects of GMFs. Educated members of the public had positive perception on the benefits and use of GMFs. There is need for public education and sensitization on GMFs to dispel misinformation and increase on the acceptability of GMFs. There is need for the policy makers and the government of Uganda to increase public education and awareness on GMFs. I believe that public trust may be an important tool that should be strengthened to bridge the gap between the public, scientists and the government hence need to include a section in this bill that that mandates strict liability and redress against any harm caused as by any research. Finally, the government of Uganda should involve the public at all levels before the Biosafety and Biotechnology bill can be passed.

## Declarations

### Ethics approval and consent to participate

Ethical approval was sought from the School of Biomedical Sciences Higher Degrees Research and Ethics Committee and permission was obtained from area local council chairpersons. Informed consent was obtained from all participants prior to recruitment into the study.

### Author’s contribution

**MN**-contributed to the conception and designing of the study, data collection, analysis and interpretation of data and manuscript writing.

## Acknowledgments

The author would like to express her gratitude to all the participants who took part in the study

## REFERENCES

1. The world counts. People who died from hunger 2022 [Available from: https://www.theworldcounts.com/challenges/people-and-poverty/hunger-and-obesity/how-many-people-die-from-hunger-each-year/story.

2. Wheeler T, Von Braun J. Climate change impacts on global food security. Science. 2013;341(6145):508–13.

3. turral H, Burke J, Faurès J-M. Climate change, water and food security: Food and agriculture organization of the United nations (FAO); 2011.

4. World Bank. Beyond hunger: ensuring food security for all 2020 [Available from: https://datatopics.worldbank.org/sdgatlas/goal-2-zero-hunger/.

5. Nutrition books. Undernutrition Diseases. 2022.

6. Afzal H, Zahid K, Ali Q, Sarwar K, Shakoor S, Nasir U, et al. Role of biotechnology in improving human health. J Mol Biomark Diagn. 2016;8(309):2.

7. Lobell DB, Gourdji SM. The influence of climate change on global crop productivity. Plant physiology. 2012;160(4):1686–97.

8. Esquinas-Alcázar J. Protecting crop genetic diversity for food security: political, ethical and technical challenges. Nature Reviews Genetics. 2005;6(12):946–53.

9. Lassoued R, Hesseln H, Phillips PW, Smyth SJ. Top plant breeding techniques for improving food security: an expert Delphi survey of the opportunities and challenges. International Journal of Agricultural Resources, Governance and Ecology. 2018;14(4):321–37.

10. Vergragt PJ, Brown HS. Genetic engineering in agriculture: New approaches for risk management through sustainability reporting. Technological Forecasting and Social Change. 2008;75(6):783–98.

11. Bauer MW. Controversial medical and agri-food biotechnology: a cultivation analysis. Public understanding of science. 2002;11(2):93.

12. Lucht JM. Public acceptance of plant biotechnology and GM crops. Viruses. 2015;7(8):4254–81.

13. Chagwena D, Sithole B, Masendu R, Chikwasha V, Maponga C. Knowledge, attitudes and perceptions towards genetically modified foods in Zimbabwe. African Journal of Food, Agriculture, Nutrition and Development. 2019;19(3):14752–68.

14. Schnurr MA, Gore C. Getting to ‘yes’: Governing genetically modified crops in Uganda. Journal of international development. 2015;27(1):55–72.

15. Monitor D. Museveni declines to sign GMO bill into law. 2017.

16. Statistics UBo. The national population and housing census 2014–main report. UBOS Kampala, Uganda; 2016.

17. Frewer L, Lassen J, Kettlitz B, Scholderer J, Beekman V, Berdal KG. Societal aspects of genetically modified foods. Food and Chemical toxicology. 2004;42(7):1181–93.

18. Ekici K, Sancak YC. A perspective on genetically modified food crops. African Journal of Agricultural Research. 2011;6(7):1639–42.

19. Lähteenmäki L, Grunert K, Ueland Ø, Åström A, Arvola A, Bech-Larsen T. Acceptability of genetically modified cheese presented as real product alternative. Food quality and preference. 2002;13(7-8):523–33.

20. Rowe G. How can genetically modified foods be made publicly acceptable? TRENDS in Biotechnology. 2004;22(3):107–9.

21. Miles S, Frewer LJ. Investigating specific concerns about different food hazards. Food quality and preference. 2001;12(1):47–61.

22. Huang J, Qiu H, Bai J, Pray C. Awareness, acceptance of and willingness to buy genetically modified foods in Urban China. Appetite. 2006;46(2):144–51.

23. Februhartanty J, Widyastuti TN, Iswarawanti DN. Attitudes of agricultural scientists in Indonesia towards genetically modified foods. Asia Pacific Journal of Clinical Nutrition. 2007;16(2).

24. Buah J. Public perception of genetically modified food in Ghana. American Journal of Food Technology. 2011;6(7):541–54.

25. Kushwaha S, Musa A, Lowenberg-DeBoer J, Fulton JR. Consumer acceptance of GMO cowpeas in sub-Sahara Africa. 2004.

26. Peter L. An investigation into the consumer acceptance of genetically modified foods at the Chris Hani district municipality, Eastern Cape, South Africa. Kuwait Chapter of the Arabian Journal of Business and Management Review. 2014;3(11):264.

27. Cui K, Shoemaker SP. Public perception of genetically-modified (GM) food: a nationwide Chinese consumer study. npj Science of Food. 2018;2(1):1–8.

28. Zhang M, Chen C, Hu W, Chen L, Zhan J. Influence of source credibility on consumer acceptance of genetically modified foods in China. Sustainability. 2016;8(9):899.

29. Öz B, Unsal F, Movassaghi H. Consumer attitudes toward genetically modified food in the United States: Are Millennials different? Journal of Transnational Management. 2018;23(1):3–21.

30. thompson PB. Ethics, hunger, and the case for genetically modified (GM) crops. Ethics, Hunger and Globalization: Springer; 2007. p. 215–35.

31. Quisumbing AR, Brown LR, Feldstein HS, Haddad L, Peña C. Women: The key to food security. Food and Nutrition Bulletin. 1996;17(1):1–2.

32. Abedi T, Alemzadeh A. Public knowledge and acceptance of genetically modified organisms in Shiraz, Iran. Age. 15(18):14.

33. Purchase IF. What determines the acceptability of genetically modified food that can improve human nutrition? Toxicology and Applied Pharmacology. 2005;207(2):19–27.

34. Bonah E, Issah NG, Kunyangna P. Consumer Knowledge, Perceptions and Acceptance of Genetically Modified Foods among Residents in the Tamale Metropolis, Ghana. American Journal of Food Science and Nutrition Research. 2017;4(3):87–98.

35. Mnaranara T, Zhang J, Wang G. Public perception towards genetically modified foods in Tanzania. Journal of Animal & Plant Sciences. 2017;27(2):586–602.

36. Pham N, Mandel N. What influences consumer evaluation of genetically modified foods? Journal of Public Policy & Marketing. 2019;38(2):263–79.

37. Muzhinji N, Ntuli V. Genetically modified organisms and food security in Southern Africa: conundrum and discourse. GM Crops & Food. 2021;12(1):25–35.

38. De Beer T, Wynberg R. Developing and implementing policy for the mandatory labelling of genetically modified food in South Africa. South African Journal of Science. 2018;114(7-8):s98–104.

